# StaRFM: Confidence-Aware Adaptation for Medical Image Segmentation Under Acquisition Shift

**DOI:** 10.64898/2026.11.22.768403

**Authors:** Behraj Khan, Tahir Qasim Syed, Shabir Ahmad, Syed Ahmad Chan Bukhari

**Author notes:** Corresponding authors: Syed Ahmad Chan Bukhari. Behraj Khan and Tahir Qasim Syed contributed equally to this work and share first authorship. (B. Khan).

## Abstract

Medical image segmentation models can lose accuracy and produce unreliable confidence estimates when deployed across acquisition centres. We study StaRFM, an adaptation objective that combines a Fisher information penalty (FIP) with a confidence misalignment penalty (CMP), and examine how the spatial aggregation of CMP affects lesion segmentation. We evaluate a Segment Anything Model (SAM) adaptation on five MRI and CT datasets and conduct a controlled leave-centre-out experiment on ISLES 2022 using a 3D U-Net. The SAM experiments improve mean Dice by 2.3 percentage points relative to fine-tuned SAM; calibration was not measured for those models. In the ISLES experiment, the baseline achieved 73.0% Dice, 7.71% boundary expected calibration error (bECE), and 1.03 ×10^−2^ negative log-likelihood (NLL) on 52 held-out-centre cases. A class-balanced CMP variant achieved 74.8% Dice, 6.57% bECE, and 0.59 × 10^−2^ NLL; a boundary-restricted variant achieved 74.1% Dice, 6.24% bECE, and 0.58 × 10^−2^ NLL. In contrast, stronger voxel-averaged CMP reduced Dice, and FIP offered no measurable additional benefit in this controlled study. Because the balanced and boundary variants were developed after an initial negative experiment, these results are exploratory. Complementary vision-language experiments and mathematical analysis characterise the broader objective, without establishing clinical reliability or a general calibration guarantee under shift. Our findings highlight the importance of class imbalance and boundary-aware evaluation when designing confidence penalties for medical segmentation.

## 1. Introduction

Large pre-trained models have changed how image analysis systems are built. Contrastive vision-language models such as CLIP Radford, Kim, Hallacy, Ramesh, Goh, Agarwal, Sastry, Askell, Mishkin, Clark et al. (2021) learn transferable image-text representations from web-scale data, and promptable segmentation models such as the Segment Anything Model (SAM) Kirillov, Mintun, Ravi, Mao, Rolland, Gustafson, Xiao, Whitehead, Berg, Lo et al. (2023) segment new objects from a box or point prompt. Medical variants followed quickly. MedSAM fine-tunes SAM on more than a million medical image-mask pairs Ma, He, Li, Han, You and Wang (2024), SAM-Med3D re-trains a fully volumetric promptable model Wang, Guo, Ye, Deng, Cheng, Li, Chen, Su, Huang, Shen, Fu, Zhang, He and Qiao (2024a), Med-SA inserts lightweight adapters into SAM Wu, Wang, Hong, Ji, Fu, Xu, Xu and Jin (2025), and BiomedCLIP Zhang, Xu, Usuyama, Xu, Bagga, Tinn, Preston, Rao, Wei, Valluri et al. (2023) and CXR-CLIP You, Gu, Ham, Park, Kim, Hong, Baek and Roh (2023) specialise contrastive pre-training to biomedical figures and chest radiographs. Systematic evaluations show that SAM’s out-of-the-box accuracy on medical images varies widely with the task, the anatomy and the prompt Mazurowski, Dong, Gu, Yang, Konz and Zhang (2023); Huang, Yang, Liu, Zhou, Chang, Zhou, Chen, Yu, Chen, Chen et al. (2024). In practice these models are therefore adapted with a small number of labelled cases from the target task.

Two problems limit what such adapted models can safely be used for. The first is *distribution shift*: the data seen at deployment differ from those used for adaptation, *P*_tr_(x, y) ≠| *P*_ts_(x, y). A useful modelling assumption is *covariate shift*, in which the image distribution changes with scanner vendor, field strength, protocol or reconstruction while the conditional label distribution *P* (y | x) remains fixed Shimodaira (2000). Actual cross-centre differences may also involve lesion prevalence, annotation conventions or case mix, so this assumption is not guaranteed. The second is *miscalibration*: predicted probabilities do not match empirical frequencies Guo, Pleiss, Sun and Weinberger (2017). The two interact. Large benchmarks show that calibration degrades as shift grows Ovadia, Fertig, Ren, Nado, Sculley, Nowozin, Dillon, Lakshminarayanan and Snoek (2019). Segmentation networks trained with Dice-type losses are systematically overconfident even in distribution Mehrtash, Wells, Tempany, Abolmaesumi and Kapur (2020); Yeung, Rundo, Nan, Sala, Schönlieb and Yang (2023), and popular CLIP adaptation methods lose much of the zero-shot model’s calibration under distribution drift Murugesan, Silva-Rodríguez, Ayed and Dolz (2025). For a lesion segmentation model, overconfidence is most harmful where it is most likely to occur: at ambiguous lesion boundaries and on images from an unfamiliar scanner.

Existing remedies tend to address one of these problems at a time. Domain generalisation methods train on source data alone and try to anticipate acquisition changes, for example with randomised intensity and spatial augmentations Ouyang, Chen, Li, Li, Qin, Bai and Rueckert (2023). Test-time adaptation updates a normalisation module or the network itself on each unlabelled test image or batch Karani, Erdil, Chaitanya and Konukoglu (2021); Wang, Shelhamer, Liu, Olshausen and Darrell (2021), and source-free adaptation removes the need for source data Hu, Liao and Xia (2026). These methods target accuracy and rarely report calibration. Conversely, a growing family of train-time calibration losses for segmentation smooths labels in a spatially varying way Islam and Glocker (2021), maximises entropy on erroneous pixels Larrazabal, Martínez, Dolz and Ferrante (2023), constrains logit margins Murugesan, Liu, Galdran, Ben Ayed and Dolz (2023b); Murugesan, Adiga Vasudeva, Liu, Lombaert, Ben Ayed and Dolz (2023a), or directly minimises a differentiable calibration error Barfoot, Garcia-Peraza-Herrera, Glocker and Vercauteren (2024). These losses are usually trained and evaluated within one domain, with few exceptions Matzkin, Larrazabal, Milone, Dolz and Ferrante (2025), and they do not come with guarantees under shift.

In earlier work we proposed CalShift Khan, Qureshi, Durrani and Syed (2025), which combines a Fisher information penalty (FIP) with a confidence misalignment penalty (CMP) for 2D image classification. A preliminary extension to SAM for brain MRI segmentation appeared as a preprint Khan and Syed (2025b). Moving these ideas to volumetric foundation-model adaptation raises three questions that the earlier work did not settle. How should the penalties be computed for 3D volumes, where a full Fisher matrix is out of reach? What, precisely, does each penalty optimise at the voxel level? And which guarantees actually hold?

This paper answers these questions with **StaRFM** (Stable and Robust Foundation Model adaptation), a single training objective that adds FIP and CMP to the loss used for CLIP prompt learning or SAM fine-tuning. FIP is computed patch-wise (16^3^ voxels) from per-voxel scores, so its cost grows with the number of sampled patches rather than with volume size. CMP is defined voxel-wise. We show that it is exactly an exponential loss on the logit margin, which clarifies both why it suppresses confidently wrong voxels and why it need not reduce calibration error. On the theoretical side, we replace heuristic arguments with two statements that hold without approximation. The first is a covariate-shift transfer bound for PAC-Bayesian predictors. The second is a deterministic chain linking voxel ECE, the Brier score and CMP. The penalties are switched on only when at least one labelled adaptation case is available; with no labels (*λ*_1_ = *λ*_2_ = 0) StaRFM reduces to the pre-trained model.

We evaluate StaRFM on eleven vision-language benchmarks, eight covariate-shift benchmarks and five MRI and CT segmentation datasets. We complement these with a controlled leave-centre-out study on ISLES 2022 Hernandez Petzsche, de la Rosa, Hanning, Wiest, Valenzuela, Reyes, Meyer, Liew, Kofler, Ezhov et al. (2022). That study fixes a single architecture and training budget, holds out an entire acquisition centre, and reports a full calibration panel with patient-level statistics. Its results show that how CMP is averaged over a volume matters. Averaged over all voxels, most of which are background, CMP lowered Dice on the held-out centre. Balanced between lesion and background, it improved boundary ECE, NLL and the Brier score on both test sets without harming Dice. FIP added no measurable benefit in this setting. We report both rounds of the study in full.

## Contributions

1. **One objective for 2D and 3D adaptation**. StaRFM adds FIP and CMP to CLIP prompt learning and to SAM fine-tuning without changing either architecture. For volumes, FIP is the patch-averaged trace of the empirical Fisher information, and CMP is applied voxel-wise.
2. **What the penalties optimise**. We show that voxelwise CMP is an exponential loss on the logit margin (Lemma 3.1). We also show that, at a sigmoid output head, the empirical Fisher trace is a feature-weighted squared error (Remark 3.1). These identities explain the behaviour we observe experimentally.
3. **Guarantees that hold as stated**. Under covariate shift, the target risk of a stochastic predictor is bounded by its source PAC-Bayes bound inflated by an importance-weight factor that no regulariser can reduce (Proposition 3.1). For binary voxel predictions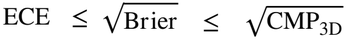 for any binning (Proposition 3.2). The role of FIP in the curvature term is given as a second-order approximation and labelled as such.
4. **Evaluation**. We report results on eleven vision-language, eight covariate-shift and five medical segmentation benchmarks (BraTS, ATLAS, ISLES 2022, Decathlon Liver and Pancreas). We add a leave-centre-out ISLES 2022 study with Dice, HD95, pooled and boundary ECE, Brier score, negative log-likelihood, patient-level boot-strap confidence intervals, Wilcoxon tests and Holm correction. The study compares voxel-wise, class-balanced and boundary forms of CMP and reports a first round in which the voxel-wise form lowered Dice.

Figure 1 shows the workflow for vision-language adaptation and Figure 2 the workflow for medical segmentation.

**Figure 1:**
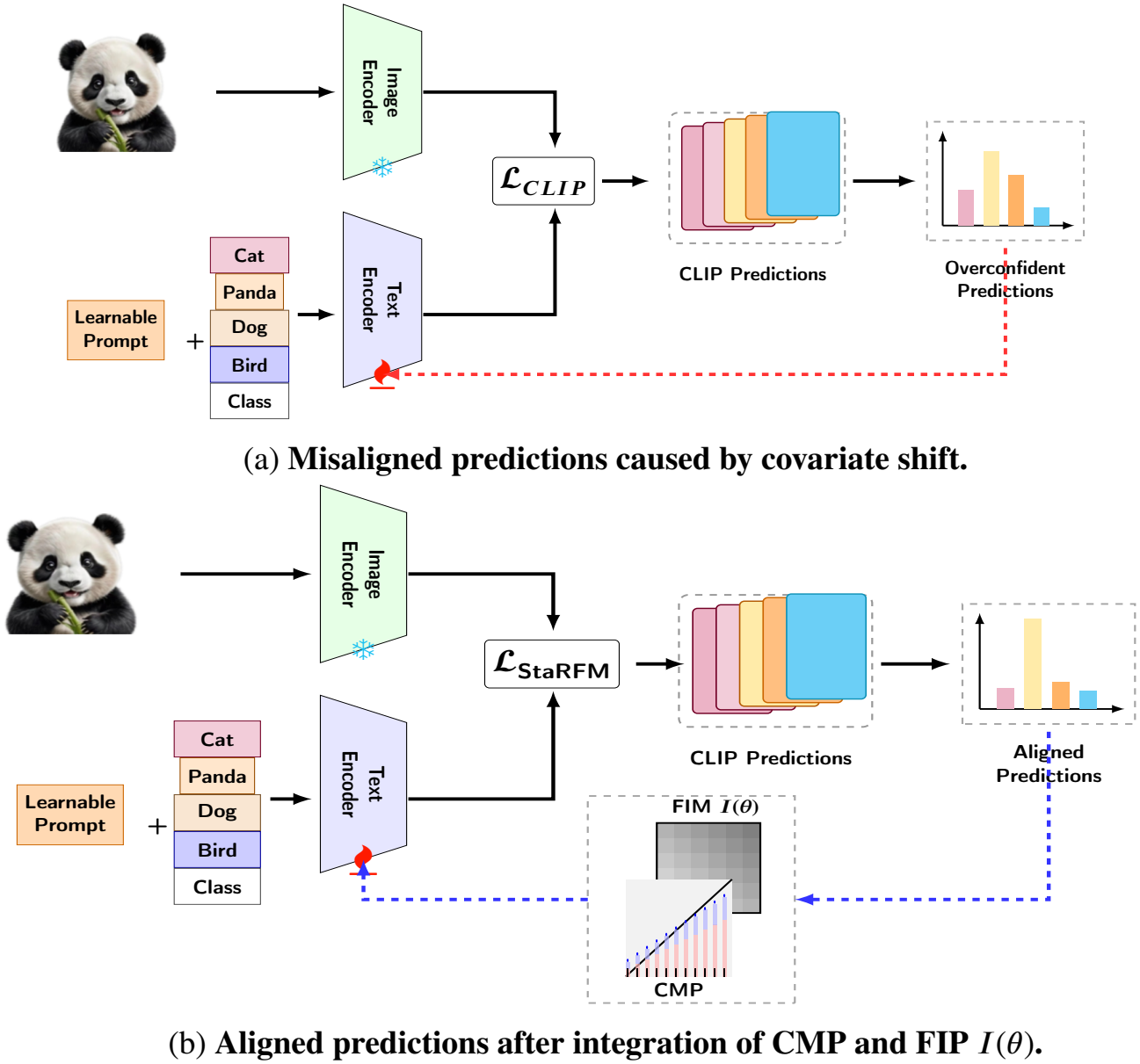
Workflow of the proposed StaRFM framework. (a) Confidence misalignment caused by covariate shift; overconfident predictions are shown bottom-right. (b) FIP (*I*(*θ*)) and CMP are integrated into the CLIP **text encoder** (the fire icon denotes the trainable component; the snowflake icon denotes a frozen encoder). The final lossℒ_StaRFM =_ ℒ_SAM_ + *λ*_1_ *I*(*θ*) + *λ*_2_ CMP produces aligned predictions (bottom-right).

**Figure 2:**
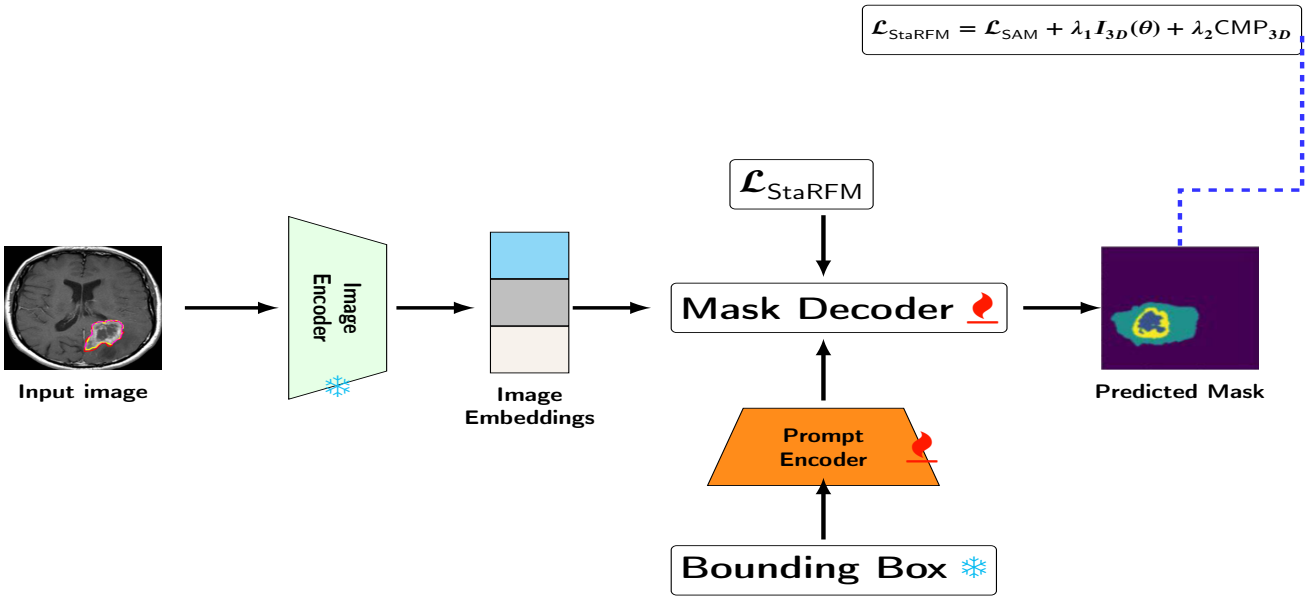
Workflow of StaRFM for medical segmentation. FIP and CMP are integrated into the SAM **prompt encoder** 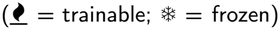. The mask decoder is also fine-tuned end-to-end. The final loss ℒ_StaRFM =_ ℒ_SAM_+ *λ*_1_*I*_3*D*_(*θ*) + *λ*_2_CMP_3*D*_combines the SAM loss with FIP and CMP.

## 2. Related Work

### 2.1. Foundation models for medical image segmentation

SAM Kirillov et al. (2023) was trained on over one billion masks and segments objects from box, point or mask prompts. Its transfer to medical images is uneven. Mazurowski et al. Mazurowski et al. (2023) found that single-prompt accuracy depends strongly on the dataset and task, and is better with box prompts than with point prompts and for well-delineated organs than for tumours. Huang et al. Huang et al. (2024) report similar variability across a large collection of medical tasks. Three adaptation strategies dominate. Full or decoder-only fine-tuning on large medical corpora is used by MedSAM Ma et al. (2024). Parameter-efficient adapters are used by Med-SA Wu et al. (2025). Native 3D re-training is used by SAM-Med3D Wang et al. (2024a), which addresses the slice-wise 2D design of the original model. On the vision-language side, Biomed-CLIP Zhang et al. (2023) and CXR-CLIP You et al. (2023) adapt contrastive pre-training to biomedical data. Prompt learning for medical VLMs combines visual and textual prompts Wang, Zhang and Yuan (2024b) or injects graph-structured knowledge for few-shot recognition Xie, Qiu, Wang, Chen, Fang, Tang and Zhang (2026). These works aim at accuracy. Calibration and shift robustness of the adapted model are usually not reported, which is the gap StaRFM addresses.

### 2.2. Distribution shift in medical image segmentation

Acquisition differences between sites are a well-documented cause of performance loss in segmentation networks Karani et al. (2021). Three families of methods address the problem. *Domain generalisation* trains on one or more source domains and anticipates unseen ones. Ouyang et al. Ouyang et al. (2023), for example, synthesise intensity and texture shifts with randomly weighted shallow networks. *Test-time adaptation* updates part of the model on each test image or batch. Karani et al. Karani et al. (2021) adapt a shallow normalisation network under a learned shape prior, and TENT Wang et al. (2021) minimises prediction entropy by updating the affine parameters of normalisation layers; related online schemes adapt on streams of unlabelled test batches Chen, Xu, Xu, Zhang and Cui (2024). *Unsupervised and source-free adaptation* aligns source and target features with or without access to source data Ganin, Ustinova, Ajakan, Germain, Larochelle, Laviolette, March and Lempitsky (2016). ProSFDA combines domain-aware image prompts with feature alignment Hu et al. (2026), and UM-SAM filters SAM-generated pseudo-labels and distils them into a segmentation network Fu, Li, Lu, Zhang and Wang (2025). All three families target accuracy when target labels are scarce or absent. StaRFM addresses a different regime: a few labelled adaptation cases are available, and the objective must improve robustness and calibration together. TTA methods are complementary and could be applied on top of a StaRFM-adapted model; we do not evaluate that combination here.

### 2.3. Calibration of segmentation networks

Mehrtash et al. Mehrtash et al. (2020) showed that segmentation networks trained with Dice loss are poorly calibrated and that ensembling helps. Jungo and Reyes Jungo and Reyes (2019) found that voxel-wise uncertainty estimates can be well calibrated on average yet unreliable for individual subjects. Yeung et al. Yeung et al. (2023) identified the Dice loss itself as a source of overconfidence and proposed DSC++, a variant that penalises overconfident predictions more strongly. Train-time calibration losses form the closest body of work to CMP:

- Spatially varying label smoothing (SVLS) blurs the one-hot target with a spatial kernel so that uncertainty concentrates at boundaries Islam and Glocker (2021).
- MEEP maximises the entropy of the predictive distribution only on misclassified pixels Larrazabal et al. (2023).
- Margin-based label smoothing (MbLS) recasts label smoothing and focal loss as equality constraints on logit distances and relaxes them to inequality constraints Murugesan et al. (2023b). NACL extends the constraint to neighbourhood class proportions Murugesan et al. (2023a).
- Boundary-weighted logit consistency regularises logits across stochastic transformations, with more weight near boundaries Karani, Dey and Golland (2023).
- DOMINO uses semantic class confusability Stolte, Volle, Indahlastari, Albizu, Woods, Brink, Hale and Fang (2022).
- mL1-ACE directly minimises a differentiable average calibration error Barfoot et al. (2024).

Like MEEP, CMP acts most strongly on confidently wrong voxels. Unlike MEEP, it is an exponential function of the logit margin and also acts, with exponentially decaying weight, on correctly classified voxels (Lemma 3.1). Most of these losses are evaluated within a single domain. Matzkin et al. Matzkin et al. (2025) study maximum-entropy regularisation for white-matter hyperintensity segmentation under scanner shift. They report better calibration and a stronger link between uncertainty and error, which motivates evaluating calibration explicitly on held-out acquisition sites, as we do in Section 5.4. Probabilistic segmentation models Kohl, Romera-Paredes, Meyer, De Fauw, Ledsam, Maier-Hein, Eslami, Jimenez Rezende and Ronneberger (2018) capture multi-modal label uncertainty but require dedicated architectures. Recent guidelines recommend reporting calibration alongside overlap and boundary metrics, with the metric chosen to fit the clinical question Maier-Hein, Reinke, Godau, Tizabi et al. (2024).

#### Relation to existing medical calibration studies

Karimi and Gholipour Karimi and Gholipour (2022) examine calibration and out-of-distribution detection in medical segmentation through multi-task learning and feature analysis. Their setting differs from our explicit train-time confidence penalty and leave-centre-out comparison. Prior boundary-weighted logit-consistency regularisation Karani et al. (2023) likewise shows why calibration near ambiguous segmentation boundaries deserves separate evaluation. Our contribution is not the first use of spatially weighted calibration regularisation; it is the controlled comparison of voxel-averaged, class-balanced and boundary-restricted CMP within the StaRFM framework. In particular, the latter two variants remain exploratory because they were developed after observing the initial voxel-wise result.

### 2.4. Calibration of adapted vision-language models

Prompt learning Zhou, Yang, Loy and Liu (2022) adapts CLIP by optimising context vectors while keeping both encoders frozen. Extensions condition the context on the image, couple vision and language prompts, or regularise towards hand-crafted prompts. Several studies show that adaptation harms calibration. Murugesan et al. Murugesan et al. (2025) trace the degradation under distribution drift to growing logit ranges and rescale logits towards the zeroshot range. C-TPT relates calibration to the dispersion of text features during test-time prompt tuning Yoon, Yoon, Tee, Hasegawa-Johnson, Li and Yoo (2024). DAC scales the temperature according to the distance between novel and base classes Wang, Wang, Wang, Zhang, Zhou and Wei (2024c). These methods are post-hoc or test-time corrections. StaRFM instead adds penalties to the adaptation objective itself and uses the same penalties for segmentation.

### 2.5. Fisher information and generalisation bounds

The Fisher information matrix measures how sensitive the likelihood is to parameter perturbations. It has been used to protect important weights in continual learning Kirkpatrick, Pascanu, Rabinowitz, Veness, Desjardins, Rusu, Milan, Quan, Ramalho, Grabska-Barwinska, Hassabis, Clopath, Kumaran and Hadsell (2017) and to define the Fisher-Rao norm, an invariant capacity measure Liang, Poggio, Rakhlin and Stokes (2019). Jastrzebski et al. Jastrzebski, Arpit, Astrand, Kerg, Wang, Xiong, Socher, Cho and Geras (2021) showed that a large Fisher trace early in training goes with poor generalisation, and that penalising the trace explicitly improves it. Kunstner et al. Kunstner, Hennig and Balles (2019) caution that the *empirical* Fisher, built from observed rather than model-sampled labels, does not in general capture curvature. We use the empirical Fisher and state where this matters. PAC-Bayes bounds McAllester (1999) control the risk of a randomised predictor through its empirical risk and a KL divergence to a prior. Dziugaite and Roy Dziugaite and Roy (2017) made such bounds non-vacuous for deep networks by optimising a Gaussian posterior around the trained weights, which connects them to flat minima. Under covariate shift, importance weighting relates target to source risk, and its learning guarantees depend on the second moment of the weights, a Rényi divergence between the two distributions Cortes, Mansour and Mohri (2010). Proposition 3.1 combines these two ingredients. Our earlier analysis of Fisher information under covariate shift Khan, Mirza and Syed (2024); Khan and Syed (2025a) was restricted to 2D classification.

### 2.6. Positioning

Table 1 summarises how StaRFM relates to the methods we compare against experimentally.

**Table 1:** Properties of StaRFM and the methods it is compared with. “Shift”: explicit mechanism for distribution shift; “Calib.”: a calibration objective; “3D”: native volumetric processing; “Analysis”: formal statements about risk or calibration.

| Method | Shift | Calib. | 3D | Analysis |
|---|---|---|---|---|
| CoOp Zhou et al. (2022) | ✗ | ✗ | ✗ | ✗ |
| MedSAM Ma et al. (2024) | ✗ | ✗ | ✗ | ✗ |
| SAM-Med3D Wang et al. (2024a) | ✗ | ✗ | ✓ | ✗ |
| *C*<sup>3</sup>-CLIP Khan et al. (2024) | ✓ | ✗ | ✗ | ✗ |
| CalShift Khan et al. (2025) | ✓ | ✓ | ✗ | ✓ |
| StaRFM (ours) | ✓ | ✓ | ✓ | ✓ |

## 3. Method

StaRFM adds two penalties to the loss used to adapt a pre-trained model. We first fix notation, then define the penalties for classification and for voxel-wise segmentation, state the resulting objectives, and analyse them in Section 3.5.

### 3.1. Problem setting

Let *P*_s_ and *P*_t_ denote the source (pre-training or development) and target (deployment) distributions over inputs and labels. Under covariate shift the marginals differ, *P*_s_(x) ≠ *P*_t_(x), while *P*_s_(y | x) = *P*_t_(y | x). Adaptation uses a small labelled set 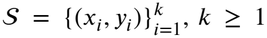.When *K* = 0 both penalties are switched off (*λ*_1_ = *λ*_2_ = 0) and the pretrained model is used unchanged; the 0-shot rows of Table 2 correspond to this case.

**Table 2:** Few-shot results on ImageNet. Top: top-1 accuracy (%) with and without the Fisher information penalty (FIP); higher is better. Bottom: calibration (ECE %) with and without the confidence misalignment penalty (CMP); lower is better.

| Method | 1-shot | 2-shot | 4-shot | 8-shot | 16-shot | Avg | Δ | Rank |
|---|---|---|---|---|---|---|---|---|
| *Accuracy (%)* |  |  |  |  |  |  |  |  |
| CoOp | 66.3 | 68.5 | 70.1 | 71.8 | 72.4 | 69.8 | – | 2 |
| CoOp + FIP | 68.1 | 70.4 | 72.0 | 73.6 | 74.3 | 71.7 | +1.9 | 1 |
| *ECE (%)* |  |  |  |  |  |  |  |  |
| CoOp | 4.82 | 4.31 | 3.88 | 3.52 | 3.36 | 3.98 | – | 2 |
| CoOp + CMP | 4.15 | 3.74 | 3.39 | 3.12 | 3.06 | 3.49 | −0.49 | 1 |

#### Vision-language classification

Given an image x ∈ ℝ^*H*×*W* ×3^ and class prompts 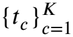, CLIP produces class probabilities *P*_*θ*_(y|x) ∈ Δ^K−1^ by a softmax over scaled cosine similarities. Following CoOp Zhou et al. (2022), both encoders are frozen and the trainable parameters *θ* are the learnable context vectors of the prompts.

#### Medical segmentation

Given an image volume x with voxel set *V*, the network outputs a logit *s* and a foreground probability *P*_*v*_ = *σ*(*s*_*v*_) for every *v* ∈ *V*. The reference mask is y_*v*_ ∈ {0, 1}. With SAM, the image encoder is frozen and the prompt encoder and mask decoder are trained (Figure 2).

### 3.2. Fisher information penalty (FIP)

For a conditional model *P*_*θ*_(y | x), the Fisher information is

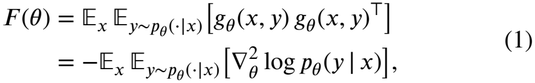

where *g*_*θ*_(x, y) = *∇*_*θ*_ log *P*_*θ*_(y x) is the score and the second equality is the information-matrix identity. In practice we use the *empirical* Fisher 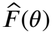, which replaces the inner expectation by the observed label, and penalise its trace. Its trace is the mean squared norm of the per-sample score:

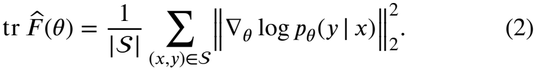

The trace is the sum of the diagonal entries, so no matrix is ever formed. The empirical Fisher coincides with *F* only near a well-specified optimum Kunstner et al. (2019); we return to this point in Remark 3.1.

#### Vision tasks

FIP is computed over the image-text pairs of the adaptation set,

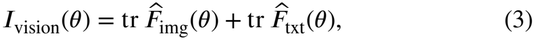

where the two terms collect the score with respect to the image-side and text-side parameters. Because both encoders are frozen, the penalty acts on training only through its dependence on the learnable context vectors.

#### Medical tasks

For volumes, the log-likelihood factorises over voxels, log *p*_*θ*_(*y* | *x*) = ∑_*v*_ log *p*_*θ*_(*y*_*v*_ | *x*). We sample a set *P* of 16^3^ patches and average the per-voxel empirical Fisher trace within each patch:

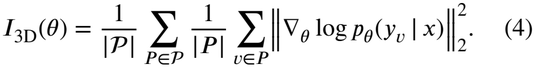

The cost depends on the number and size of the sampled patches rather than on the volume size.

##### Remark 3.1

(FIP at a sigmoid output head). Let the output layer be a 1×1×1 convolution, *s*_*v*_ = *w*^⊤^*h*_*v*_ + *b*, with input features *h*_*v*_. Then ∇(*w,b*) log *p*_*θ*_(*y*_*v*_| *x*) = (*y*_*v*_ − *p*_*v*_) [*h*_*v*_; 1], so the per-voxel empirical Fisher trace with respect to the head is

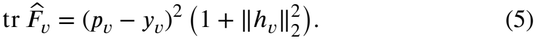

At the output head, FIP is therefore a squared-error (Brier-type) term weighted by the feature norm. Gradients flow through both *p*_*v*_ and *h*_*v*_ into the rest of the network. The *expected* Fisher replaces (*p*_*v*_ − *y*_*v*_)^2^ by *p*_*v*_(1 − *p*_*v*_), which is minimised by confident predictions whatever the label. We therefore use the empirical form whenever FIP is meant to support calibration.

### 3.3 Confidence misalignment penalty (CMP)

For *K*-class prediction with true label *y*, CMP penalises every wrong class that is ranked above the true class:

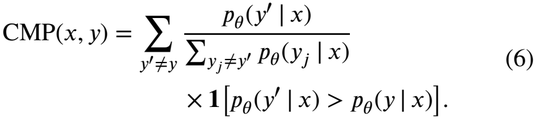

For vision tasks, CMP_vision_ is the mean of Eq. (6) over the adaptation set.

#### Medical tasks (voxel-wise)

In the binary case the denominator of Eq. (6) is 1 − *p*_*θ*_(*y*^′^ | *x*) with 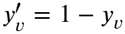. We drop the indicator to obtain a smooth surrogate. Each term is non-negative, so the surrogate upper-bounds the indicator version. Let *q*_*v*_ = |*p*_*v*_ −*y*_*v*_ |be the probability assigned to the wrong label, and let 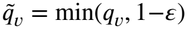 for a small 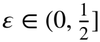.

Then

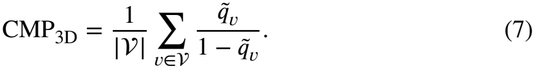

##### Lemma 3.1

(CMP_3D_ is an exponential margin loss). *Let m*_*v*_ = (2*y*_*v*_ − 1) *s*_*v*_ *be the signed logit margin of voxel v*.

*If q* ≤ 1 − *ε, then* 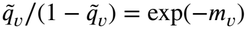.

*Proof*. For *y*_*v*_ = 1, *q*_*v*_ = 1 − *σ*(*s*_*v*_) = *σ*(−*s*_*v*_), and 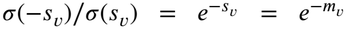. The case *y*_*v*_ = 0 is symmetric. □

Lemma 3.1 makes the behaviour of CMP_3D_ explicit, with three consequences.

- A confidently wrong voxel (*m*_*v*_ ≪ 0) contributes a term that grows exponentially in |*m*_*v*_|, until the clipping at 1 − *ε* caps it at (1 − *ε*)/*ε*. A clipped term has zero gradient, so the worst voxels would no longer be penalised. For training we therefore continue the term linearly beyond the cap, with the same value and slope at 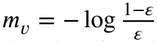. The continued term is never smaller than the clipped one. The continuation is used only as a training loss. Every reported value of CMP, and every bound below, uses the capped expression of Eq. (7).
- Correctly classified voxels (*m*_*v*_ > 0) still contribute 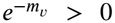 and are pushed towards larger margins. CMP therefore does not, by itself, discourage overconfidence on correct voxels.
- The population minimiser of the exponential loss is half the log-odds Friedman, Hastie and Tibshirani (2000). Used alone, CMP would thus yield probabilities that are less extreme than the true conditional probabilities. In StaRFM it is combined with BCE, which is a proper scoring rule, and with Dice, which is known to encourage overconfidence Mehrtash et al. (2020); Yeung et al. (2023). The net effect on calibration is therefore an empirical question, which Section 5.4 examines.

#### Class-balanced and boundary-restricted variants

In a lesion volume almost all voxels are background, so the mean in Eq. (7) is dominated by background terms. We therefore also consider two variants. Let *F*= {*v* : *y*_*v*_ = 1}, *B*=*V*\*F*, let *Nk* be the voxels within *k* voxels of the reference boundary (dilation minus erosion), and let 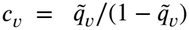. Then

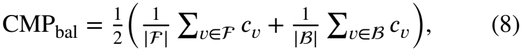

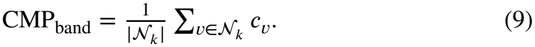

For a volume without lesion, CMP_bal_ reduces to the background mean and CMP_band_ is set to zero. Unless stated otherwise, CMP_3D_ denotes the voxel-wise form of Eq. (7). The three forms are compared in Section 5.4.

### 3.4. Training objectives

#### Vision-language

With the symmetric contrastive loss

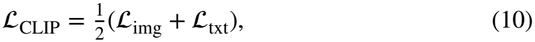

where

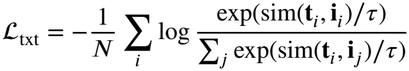

and ℒ_img_ is defined symmetrically, the objective is

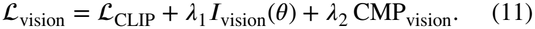

#### Medical segmentation

With soft Dice and binary cross-entropy,

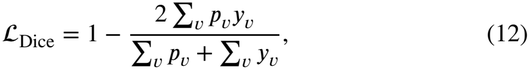

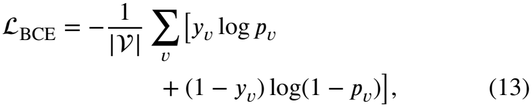

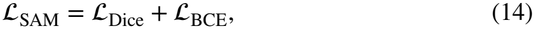

the objective is

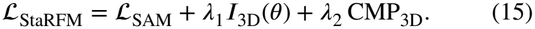

The same objective, with ℒ_SAM_ replaced by the Dice-plus-BCE loss of a U-Net, is used in Section 5.4.

### 3.5.Theoretical analysis

We state two results that hold exactly and one approximation. Proofs are in Appendix A.

#### Proposition 3.1

(Covariate-shift transfer of a PAC-Bayes bound). *Let l*(*θ*; *x, y*) ∈ [0, 1] *be a bounded loss, and let R*_*s*_(*θ*) *and R*_*t*_(*θ*) *be its source and target risks. Assume covariate shift with P*_*t*_(*x*) *absolutely continuous with respect to P*_*s*_(*x*), *importance weight w*(*x*) = d*P*_*t*_/d*P*_*s*_(*x*), *and* 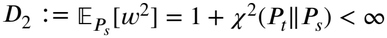.

i. *For every θ*, 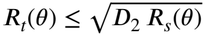.
ii. *Let* Π *be a prior over parameters chosen before seeing an i.i.d. source sample of size n, and let B*(KL, *n, δ*) *be any PAC-Bayes complexity term such that, with probability at least* 1−*δ*, 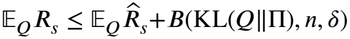 *for all posteriors Q (e.g. McAllester (1999)). Then, with the same probability, for all Q*,

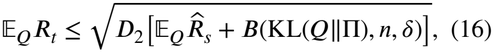 *where* E_*Q*_ *denotes expectation over θ* ∼ *Q*.

Two points about Proposition 3.1 matter for its interpretation. First, *D*_2_ depends only on the data distributions. No regulariser, FIP included, can reduce it. If the target contains image appearances that the source never produces, *D*_2_ is infinite and the bound is void. Second, FIP enters only through the source-side terms, as the following approximation shows.

#### Remark 3.2

(FIP and the curvature term; approximation). Take 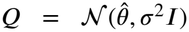 and Π =*N* (*θ*_0_, *σ*^2^*I*), so that 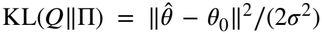. For a twice differentiable surrogate risk 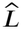(here the log-loss), 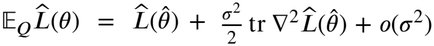. For the log-loss, the Hessian equals the Fisher in expectation over labels drawn from the model (Eq. (1)). Penalising tr 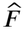 therefore targets the curvature term of the Gibbs risk in Eq. (16), in line with the flat-minimum reading of PAC-Bayes bounds Dziugaite and Roy (2017) and with the empirical findings of Jastrzebski et al. (2021). This argument is second-order and uses the surrogate rather than the bounded loss. With the empirical Fisher it holds only approximately Kunstner et al. (2019). We do not claim it as a guarantee.

#### Proposition 3.2

(Voxel calibration is controlled by Brier score and CMP). *Consider binary voxel predictions p*_*v*_ ∈ [0, 1] *with labels y*_*v*_ ∈ {0, 1}, *v* = 1, …, *N. Define the predicted label* 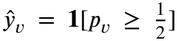, *the top-label confidence c*_*v*_ = max(*p*_*v*_, 1 − *p*_*v*_) *and the correctness* 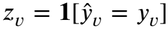. *Let B*1, …, *B*_*M*_ *be any partition of the voxels (in particular equal-width or equal-mass binning on c*_*v*_*). Let* 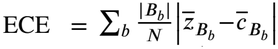, *where bars denote within-bin means, and let* 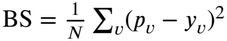. *Then, for every* 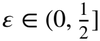,

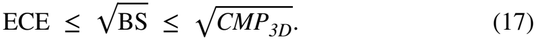

The inequalities are deterministic. They hold for every volume and for pooled voxels, with no sample-size term. An additive bound of the form ECE ≤ BS + *o*(1) cannot hold. If all labels are 0 and *p*_*v*_ = 0.1 for every voxel, then ECE = 0.1 while BS = 0.01. The square root in Eq. (17) is therefore necessary, and in this example the first inequality is tight. The chain also shows that lowering CMP or the Brier score tightens an upper bound on ECE without necessarily lowering ECE itself.

#### Corollary 3.1.1.

*If a volume contains both lesion and background voxels, then* 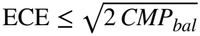.

*Proof*. With *w* = | *F*|/|*V* | and the class means *A* and *B* of c*v*, CMP3D = *wA* + (1 − *w*)*B* ≤ *A* + *B* = 2 CMP_bal_. Apply Proposition 3.2. □

No analogous bound holds for CMP_band_, which ignores voxels far from the boundary.

## 4. Experiments

The experiments test three claims. (i) FIP improves robustness to covariate shift in 2D classification and 3D segmentation. (ii) CMP improves calibration in classification.

For segmentation, we test what it does to the calibration metrics that Proposition 3.2 bounds. (iii) The objective transfers across modalities (MRI, CT) and organs (brain, liver, pancreas).

### 4.1. Datasets

#### Vision-language

We use 11 recognition datasets and 8 covariate-shift benchmarks.

- *Recognition*: ImageNet Russakovsky, Deng, Su, Krause, Satheesh, Ma, Huang, Karpathy, Khosla, Bernstein et al. (2015), Caltech101, OxfordPets, Flowers102, Food101, SUN397, DTD, EuroSAT, UCF101, FGV-CAircraft and StanfordCars. They cover generic objects, fine-grained categories, scenes, textures, actions and satellite images.
- *Covariate shift*: PACS, Office-Home, VLCS, DomainNet and four ImageNet variants. ImageNet-V2 recollects the test set, ImageNet-R contains renditions such as paintings and cartoons, ImageNet-A contains natural images that standard classifiers misclassify, and ImageNet-S contains sketches.

#### Medical segmentation

We use five public datasets spanning two modalities.

- **BraTS 2023** Menze, Jakab, Bauer, Kalpathy-Cramer, Farahani, Kirby, Burren, Porz, Slotboom, Wiest et al. (2014); Baid et al. (2021): multi-parametric brain MRI (T1, T1c, T2, FLAIR) with glioma sub-region labels, acquired at many institutions with different scanners and protocols. We use 1,250 scans.
- **ATLAS v2.0** Liew, Lo, Donnelly, Zavaliangos-Petropulu, Jeong, Barisano, Hutton, Simon, Juliano, Suri et al. (2022): T1-weighted MRI with manually traced chronic stroke lesions. The release contains 1,271 scans: 655 public training scans, 300 hidden test scans (together drawn from 33 research cohorts) and a hidden generalisability set of 316 scans.
- **ISLES 2022** Hernandez Petzsche et al. (2022): multivendor MRI (DWI, ADC, FLAIR) of acute and subacute ischaemic stroke. The dataset has 400 cases, of which the 250 training cases are public. Compared with ATLAS, ISLES covers an earlier stage of stroke and a diffusion-based protocol.
- **Decathlon Liver** Antonelli, Reinke, Bakas, Farahani, Kopp-Schneider, Landman, Litjens, Menze, Ronneberger, Summers et al. (2022): contrast-enhanced CT with liver and tumour labels (131 labelled training volumes). HU window [−200, 250].
- **Decathlon Pancreas** Antonelli et al. (2022): CT with pancreas and tumour labels (281 labelled training volumes). The organ is small, its boundaries are ambiguous, and the tumour class is strongly imbalanced. HU window [−125, 275].

Each dataset brings its own sources of covariate shift: scanner vendor, field strength, contrast protocol and slice thickness.

### 4.2. Baselines

#### Vision-language

We compare with zero-shot CLIP Radford et al. (2021) and CoOp Zhou et al. (2022). Other prompt-learning methods, and our earlier CalShift Khan et al. (2025), are discussed in Section 2 but were not re-run under the present protocol.

#### Medical segmentation

- **Vanilla SAM** Kirillov et al. (2023): zero-shot with box prompts.
- **SAM-FT**: SAM with a fine-tuned mask decoder and frozen image encoder, following Ma et al. (2024).
- **MedSAM** Ma et al. (2024): SAM fine-tuned on a large medical corpus.
- **SAM-Med3D** Wang et al. (2024a): a native 3D promptable model.
- **SAM-FT + CalShift (2D)**: SAM-FT trained with the 2D CalShift penalties Khan et al. (2025).

#### Implementation details

*Vision-language:* CLIP ViT-B/16, Adam (learning rate 10^−4^, batch size 32), 50 epochs, prompt length 16 tokens as in CoOp Zhou et al. (2022), *λ*_1_ = 0.4, *λ*_2_ = 0.5. All models start from the same CLIP checkpoint, and only the standard CLIP preprocessing is applied. *Medical:* SAM ViT-H image encoder (frozen); prompt encoder and mask decoder fine-tuned; Adam (learning rate 10^−4^); *λ*_1_ = 0.3, *λ*_2_ = 0.5; 50 epochs; patch size 128^3^. CT intensities are clipped to the dataset-specific HU window and then z-score normalised; MRI volumes are z-score normalised per volume. Box prompts are derived from the reference masks as in MedSAM Ma et al. (2024). All experiments ran on one NVIDIA A100 (80 GB).

### 4.3. Evaluation metrics

#### Classification

Top-1 accuracy and expected calibration error (ECE) Guo et al. (2017); Naeini, Cooper and Hauskrecht (2015),

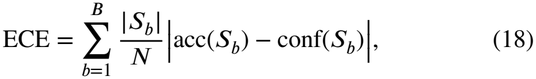

with *B* = 10 equal-width confidence bins.

#### Segmentation

Dice similarity coefficient (DSC) after thresholding at 0.5, and the 95th-percentile Hausdorff distance (HD95) in millimetres. HD95 is the larger of the two directed 95th percentiles of boundary-to-boundary distances, computed with the voxel spacing from the image header. It is undefined when either mask is empty, and such cases are excluded and counted. In the leave-centre-out study we also report voxel calibration:

- *pooled ECE*: Eq. (18) over all voxels of a volume, using the top-label confidence max(*p*_*v*_, 1 − *p*_*v*_), as in Proposition 3.2;
- *boundary ECE* (bECE): the same quantity restricted to a band around the reference boundary, obtained by three iterations of binary dilation minus three iterations of erosion with the 6-connected structuring element (voxels within city-block distance three of the boundary). Pooled ECE is dominated by easy background voxels, which is why we report bECE separately;
- the *Brier score* and the *negative log-likelihood* (NLL), both proper scoring rules;
- the predicted-to-reference lesion volume ratio, to detect systematic under- or over-segmentation;
- CMP_3D_ with *ε* = 10^−3^, in the capped form of Eq. (7).

All metrics are computed per patient and then averaged.

#### Statistics

The unit of analysis is the patient, never the voxel or the slice. For the five-dataset comparison, results are averaged over three random seeds and reported as mean±standard deviation across seeds. Significance in the medical ablation is assessed with a paired two-sided Wilcoxon signed-rank test over test patients. values, the smallest attainable two-sided p-value of this test is 0.25. Tests over seeds therefore cannot reach conventional significance levels, and we do not use them. In the leave-centre-out study we report paired per-patient differences with 95% percentile bootstrap confidence intervals (10,000 resamples of patients) and Wilcoxon p-values. The p-values are Holm-adjusted across all comparisons with the baseline and all six metrics within each evaluation split.

## 5 Results and Discussion

### 5.1. Vision-language classification

Table 3 compares StaRFM with zero-shot CLIP and CoOp on the eleven recognition benchmarks. Averaged over datasets, StaRFM raises accuracy from 71.3% to 73.1% (2.4% relative) and lowers ECE from 3.17% to 3.03% (4.5% relative). The accuracy gains are largest on Flowers102 (+6.3 points), UCF101 (+5.7) and ImageNet (+5.4). Accuracy falls on StanfordCars (−4.9) and EuroSAT (−4.6), and ECE rises on Flowers102 (2.96% to 3.16%). On five datasets (Food101, OxfordPets, ImageNet, FGVCAircraft and SUN397), zero-shot CLIP is better calibrated than both CoOp and StaRFM. This agrees with the observation that prompt learning erodes the calibration of the zero-shot model Murugesan et al. (2025). StaRFM narrows this gap but does not close it.

**Table 3:** Top-1 accuracy (%) and calibration (ECE %) on eleven vision-language benchmarks. Δ% is the relative change of StaRFM with respect to CoOp; arrows give the direction of change (for ECE, a decrease is an improvement). The last row averages over datasets, and its Δ% is computed from the averages.

|  | Accuracy (%) |  |  |  | Calibration ECE (%) |  |  |  |
|---|---|---|---|---|---|---|---|---|
| Dataset | CLIP | CoOp | StaRFM | Δ% | CLIP | CoOp | StaRFM | Δ% |
| UCF101 | 69.9 | 78.6 | 84.3 | 7.3↑ | 3.24 | 3.08 | 2.94 | 4.55↓ |
| Food101 | 90.1 | 97.0 | 98.7 | 1.7↑ | 1.57 | 3.35 | 3.02 | 9.85↓ |
| Caltech101 | 96.8 | 98.6 | 98.8 | 0.2↑ | 6.49 | 3.24 | 3.08 | 4.94↓ |
| OxfordPets | 91.2 | 98.2 | 98.9 | 0.7↑ | 2.25 | 3.06 | 2.84 | 7.19↓ |
| Flowers102 | 72.1 | 79.2 | 85.5 | 8.0↑ | 3.11 | 2.96 | 3.16 | 6.76↑ |
| ImageNet | 72.4 | 79.5 | 84.9 | 6.8↑ | 1.51 | 3.36 | 3.06 | 8.93↓ |
| StanfordCars | 63.3 | 59.2 | 54.3 | 8.3↓ | 3.74 | 3.38 | 3.16 | 6.51↓ |
| FGVCAircraft | 27.2 | 25.2 | 27.2 | 7.9↑ | 3.03 | 3.24 | 3.08 | 4.94↓ |
| SUN397 | 69.4 | 63.0 | 66.8 | 6.0↑ | 1.59 | 3.02 | 2.96 | 1.99↓ |
| DTD | 53.3 | 52.5 | 55.1 | 5.0↑ | 4.53 | 3.06 | 2.92 | 4.58↓ |
| EuroSAT | 56.5 | 53.8 | 49.2 | 8.6↓ | 9.12 | 3.08 | 3.06 | 0.65↓ |
| Average | 69.3 | 71.3 | 73.1 | 2.4↑ | 3.65 | 3.17 | 3.03 | 4.45↓ |

Table 2 reports ImageNet results from 1 to 16 shots. Adding FIP improves accuracy at every shot count (mean +1.9 points). Adding CMP lowers ECE at every shot count (mean −0.49 points), and the largest reduction occurs at one shot (4.82% to 4.15%).

### 5.2. Covariate-shift benchmarks

Table 4 reports the eight covariate-shift benchmarks. Adding FIP to CoOp improves accuracy on all eight, by 0.8 to 3.5 points. Adding CMP lowers ECE on all eight. In absolute terms the largest ECE reductions occur on ImageNet-A (0.34 points) and ImageNet-S (0.22 points), where CoOp is most overconfident. In relative terms these are the two smallest reductions (2.2% and 2.6%), and ECE remains high (15.0% and 8.2%). On ImageNet-A, ImageNet-S and ImageNet-V2, zero-shot CLIP is better calibrated than either adapted model. CMP alone is therefore far from sufficient under severe shift.

**Table 4:** Accuracy (%) and calibration (ECE %) on eight covariate-shift benchmarks. For accuracy, Δ is the absolute change of +FIP with respect to CoOp in percentage points; for ECE, Δ% is the relative reduction of +CMP with respect to CoOp.

|  | Accuracy (%) |  |  |  | Calibration ECE (%) |  |  |  |
|---|---|---|---|---|---|---|---|---|
| Dataset | CLIP | CoOp | +FIP | Δ (pp) | CLIP | CoOp | +CMP | Δ% |
| PACS | 96.1 | 96.5 | 98.0 | +1.5 | 2.18 | 2.02 | 1.91 | −5.45 |
| Office-Home | 80.4 | 82.1 | 85.6 | +3.5 | 2.77 | 2.92 | 2.75 | −5.82 |
| VLCS | 81.4 | 82.5 | 86.0 | +3.5 | 2.89 | 3.01 | 2.84 | −5.65 |
| DomainNet | 54.1 | 58.8 | 60.0 | +1.2 | 3.69 | 3.29 | 3.15 | −4.26 |
| ImageNet-V2 | 60.8 | 64.2 | 65.5 | +1.3 | 2.44 | 4.19 | 4.05 | −3.34 |
| ImageNet-S | 46.2 | 47.9 | 48.8 | +0.9 | 4.88 | 8.40 | 8.18 | −2.62 |
| ImageNet-A | 47.8 | 49.7 | 50.5 | +0.8 | 8.34 | 15.34 | 15.00 | −2.22 |
| ImageNet-R | 73.9 | 75.2 | 76.8 | +1.6 | 3.51 | 3.12 | 2.95 | −5.45 |

### 5.3. Medical image segmentation

Table 5 reports DSC on the five volumetric datasets with a SAM backbone. StaRFM obtains the highest mean DSC on every dataset, and its gains over SAM-FT range from 1.7 (Liver) to 2.7 (ISLES) points, or 2.3 points on average. The 2D CalShift penalties already recover about half of this gain (1.1 points on average over SAM-FT). The volumetric formulation adds a further 1.2 points. Averaged over the MRI datasets StaRFM reaches 80.4% DSC, and over the CT datasets 82.8%. The corresponding SAM-FT averages are 77.9% and 80.6%. For this benchmark we report DSC only. Calibration of the segmentation outputs is analysed in the controlled study below, where the full set of calibration metrics is available.

**Table 5:** Medical segmentation with a SAM backbone: DSC (%), mean ± standard deviation over three seeds. Higher is better.

| Method | BraTS | ATLAS | ISLES | Liver | Pancreas |
|---|---|---|---|---|---|
| Vanilla SAM | 78.5 ± 0.9 | 70.8 ± 1.0 | 66.2 ± 1.1 | 87.0 ± 0.7 | 62.5 ± 1.2 |
| SAM-FT | 84.0 ± 0.6 | 76.8 ± 0.7 | 72.9 ± 0.8 | 91.4 ± 0.5 | 69.8 ± 0.9 |
| MedSAM | 82.8 ± 0.7 | 75.6 ± 0.8 | 71.5 ± 0.9 | 90.5 ± 0.5 | 68.4 ± 0.9 |
| SAM-Med3D | 83.4 ± 0.6 | 76.1 ± 0.7 | 72.1 ± 0.8 | 90.9 ± 0.4 | 69.0 ± 0.8 |
| SAM-FT + CalShift (2D) | 85.0 ± 0.5 | 78.1 ± 0.6 | 74.2 ± 0.7 | 92.2 ± 0.4 | 71.0 ± 0.8 |
| StaRFM | 86.2 ± 0.4 | 79.3 ± 0.5 | 75.6 ± 0.6 | 93.1 ± 0.3 | 72.4 ± 0.7 |

### 5.4. Controlled leave-centre-out study on ISLES 2022

The SAM experiments establish accuracy gains but leave open what the penalties do to calibration under a real acquisition-site shift. We therefore ran a controlled study with three features. Every component is specified and reproducible from saved probability maps. An entire acquisition centre is held out from training and model selection. The complete calibration panel of Section 4.3 is reported with patient-level statistics. The study uses a small 3D U-Net instead of SAM so that the penalties can be studied without the confounding effects of prompting and a frozen 2D encoder. It is a single-source domain-generalisation setting: no centre-2 image is used for training, unlike the few-shot target adaptation of the main experiments.

#### Data and split

We used the 250 public ISLES 2022 training cases with per-case acquisition-centre labels: 198 cases from centre 1 and 52 from centre 2. With a fixed random seed, the centre-1 cases were split into 138 training, 30 validation and 30 in-distribution (ID) test cases. All 52 centre-2 cases form the held-out test set. The validation set was used only for checkpoint and weight selection. One ID test case has an empty reference mask. HD95 and boundary ECE are undefined for it, so their ID statistics use 29 patients.

#### Model and training

The input is the DWI and ADC volumes at native resolution, stacked as two channels, with each channel z-scored over its non-zero voxels. The network is a four-level 3D U-Net Ronneberger, Fischer and Brox (2015); Çiçek, Abdulkadir, Lienkamp, Brox and Ronneberger (2016) with 16 to 128 channels. Each level has two 3×3×3 convolutions with group normalisation and ReLU, followed by a 1×1×1 sigmoid output head (1.40 M parameters). Whole volumes are processed one at a time, zero-padded to a multiple of 8. All models minimise BCE plus soft Dice with Adam (learning rate 10^−3^) for 30 epochs. We keep the checkpoint with the highest validation DSC and threshold probabilities at 0.5. FIP is the closed-form head Fisher of Eq. (5), averaged over four random 16^3^ patches per volume. CMP uses *ε* = 10^−3^ with the linear continuation of Section 3.3. The band variant of the training loss uses *k* = 3 with a cubic (3^3^-connected) neighbourhood, which is slightly wider than the 6-connected band used to evaluate bECE. Both widths are fixed in voxels; because voxel spacing differs between scans, the physical width of the band varies from case to case. We compare a baseline, FIP alone, and, for each CMP form (voxel-wise, class-balanced, band), CMP alone and FIP plus CMP (StaRFM). Every configuration is trained with three seeds, and per-patient metrics are averaged over seeds.

#### Data integrity and reproducibility

Before training, every case was checked for matching array shapes, matching affine matrices (tolerance 10^−4^) between DWI, ADC and mask, and finite voxel values, and all input files were fingerprinted with SHA-256. Each trained model is stored with a fingerprint of its configuration, the patient split and the input-data hash, so that cached predictions from a different configuration or data version cannot be reused. Training used deterministic algorithms where PyTorch provides them (Python 3.10.12, PyTorch 2.14.1 with CUDA 13.0, NumPy 2.2.6, nibabel 5.4.2, one GPU). We re-ran the complete pipeline under this protocol, retraining all 18 weight-selection runs and all 24 final runs. Every validation score used for weight selection, and every mean, confidence interval and p-value reported below, was identical to the previous execution.

#### Weight selection

For each CMP form, (*λ*_1_, *λ*_2_) was chosen for StaRFM on the centre-1 validation set by one fixed rule: highest validation DSC after 15 epochs (seed 0), with ties within 0.001 broken by lower validation NLL. The grid was *λ*_1_ ∈ {0.001, 0.01} and *λ*_2_ ∈ {0.001, 0.01, 0.1}. The selected weights were (0.01, 0.001) for the voxel-wise form and (0.001, 0.001) for both other forms. CMP-only runs use the *λ*_2_ of the matching StaRFM run; FIP-only uses *λ*_1_ = 0.01.

The study was carried out in two rounds, and we report both. The first round used only the voxel-wise form, with a grid of *λ*_1_ ∈ {0.01, 0.1} and *λ*_2_ ∈ {0.1, 0.5}, and selected (0.01, 0.1). With these weights, StaRFM *lowered* DSC on the held-out centre from 73.0% to 66.1% (−6.9 points, 95% CI −8.2 to −5.6) and raised HD95 from 15.4 to 20.1 mm, while lowering NLL from 1.03 to 0.63 × 10^−2^. Ablation attributed the loss to CMP. The class-balanced and band variants, and the extended grid, were then defined to test whether the loss came from the background-dominated average, and were evaluated with the protocol above. Because the variants were introduced after seeing the first round, all variants are reported below, and p-values are Holm-adjusted across all seven comparisons with the baseline and all six metrics within each split (42 tests per split). This is the more conservative of the two families we computed; p-values adjusted only within each comparison (six tests) are smaller and are provided with the code.

#### Results

Table 6 lists the means and Table 7 the paired differences for the three StaRFM variants. With the selected small weight, voxel-wise StaRFM changed no metric measurably on the held-out centre, while voxel-wise CMP alone still lowered DSC by 2.1 points (95% CI −3.2 to−1.1, adjusted *p* = 0.004). The class-balanced and band forms gave consistent calibration gains on both test sets.

**Table 6:** Leave-centre-out study on ISLES 2022 (3D U-Net trained on centre 1; three seeds, 30 epochs). Means over test patients of seed-averaged per-patient metrics. Subscripts give the CMP form (Eqs. (7), (8), (9)). Weights (*λ*_1_, *λ*_2_): StaRFM (0.01, 0.001), StaRFM_bal_ and StaRFM_band_ (0.001, 0.001); single-penalty runs use the matching weight. DSC, ECE and boundary ECE (bECE) in %; HD95 in mm; Brier score ×10^−3^; NLL and CMP_3D_ ×10^−2^. HD95 and bECE on the ID split use 29 patients.

| Split | Method | DSC↑ | HD95↓ | ECE↓ | bECE↓ | Brier↓ | NLL↓ | CMP<sub>3D</sub>↓ |
|---|---|---|---|---|---|---|---|---|
| Held-out, centre 2 (*n* = 52) | Baseline | 73.0 | 15.4 | 0.119 | 7.71 | 1.25 | 1.03 | 76.3 |
|  | +FIP | 71.1 | 16.3 | 0.125 | 7.99 | 1.29 | 1.20 | 89.5 |
|  | +CMP | 70.9 | 17.3 | 0.124 | 7.88 | 1.30 | 1.13 | 83.2 |
|  | StaRFM | 72.2 | 16.8 | 0.120 | 7.63 | 1.26 | 1.03 | 79.2 |
|  | +CMP<sub>bal</sub> | 74.0 | 15.0 | 0.094 | 6.73 | 1.14 | 0.61 | 35.3 |
|  | StaRFM<sub>bal</sub> | 74.8 | 13.0 | 0.098 | 6.57 | 1.12 | 0.59 | 36.1 |
|  | +CMP<sub>band</sub> | 73.2 | 13.4 | 0.098 | 6.41 | 1.10 | 0.62 | 43.1 |
|  | StaRFM<sub>band</sub> | 74.1 | 15.1 | 0.094 | 6.24 | 1.07 | 0.58 | 38.2 |
| ID, centre 1 (*n* = 30) | Baseline | 59.6 | 30.9 | 0.124 | 8.59 | 1.28 | 1.27 | 91.8 |
|  | +FIP | 60.9 | 29.0 | 0.124 | 8.39 | 1.27 | 1.35 | 96.6 |
|  | +CMP | 60.7 | 28.4 | 0.122 | 8.54 | 1.27 | 1.25 | 93.0 |
|  | StaRFM | 60.0 | 31.8 | 0.118 | 8.16 | 1.22 | 1.13 | 86.3 |
|  | +CMP<sub>bal</sub> | 62.1 | 28.8 | 0.103 | 7.16 | 1.18 | 0.61 | 34.6 |
|  | StaRFM<sub>bal</sub> | 60.9 | 27.1 | 0.098 | 7.19 | 1.11 | 0.60 | 39.2 |
|  | +CMP<sub>band</sub> | 61.9 | 25.7 | 0.104 | 7.18 | 1.15 | 0.68 | 51.1 |
|  | StaRFM<sub>band</sub> | 59.7 | 36.0 | 0.097 | 7.11 | 1.11 | 0.61 | 42.1 |

**Table 7:** Paired per-patient differences (StaRFM variant − baseline) in the ISLES 2022 study: mean, 95% patient-level bootstrap confidence interval, and two-sided Wilcoxon p-value, Holm-adjusted across all seven comparisons with the baseline and six metrics within each split. Units as in Table 6; DSC, ECE and bECE differences in percentage points.

| Metric | StaRFM | StaRFM<sub>bal</sub> | StaRFM<sub>band</sub> |
|---|---|---|---|
| *Held-out, centre 2 (n* = 52*)* |  |  |  |
| DSC | −0.8 [−2.2, 0.4]; 1.00 | +1.8 [0.9, 2.6]; 0.001 | +1.1 [0.3, 1.8]; 0.010 |
| HD95 | +1.5 [−1.4, 4.3]; 1.00 | −2.3 [−4.7, −0.1]; 0.55 | −0.2 [−1.6, 1.2]; 1.00 |
| ECE | +0.001 [−0.003, 0.006]; 1.00 | −0.021 [−0.029, −0.015]; <0.001 | −0.026 [−0.037, −0.016]; <0.001 |
| bECE | −0.08 [−0.35, 0.26]; 0.56 | −1.13 [−1.34, −0.93]; <0.001 | −1.46 [−1.74, −1.20]; <0.001 |
| Brier | +0.01 [−0.03, 0.06]; 1.00 | −0.13 [−0.20, −0.08]; <0.001 | −0.18 [−0.26, −0.11]; <0.001 |
| NLL | +0.00 [−0.03, 0.03]; 1.00 | −0.44 [−0.67, −0.27]; <0.001 | −0.44 [−0.68, −0.27]; <0.001 |
| *ID, centre 1 (n* = 30*; HD95 and bECE n* = 29*)* |  |  |  |
| DSC | +0.4 [−5.4, 4.7]; 1.00 | +1.3 [−4.3, 5.2]; 0.16 | +0.2 [−2.8, 2.5]; 1.00 |
| HD95 | +0.9 [−1.9, 4.3]; 1.00 | −3.8 [−9.6, 1.3]; 0.77 | +5.0 [0.8, 10.0]; 1.00 |
| ECE | −0.006 [−0.012, −0.001]; 1.00 | −0.025 [−0.046, −0.011]; <0.001 | −0.027 [−0.051, −0.009]; <0.001 |
| bECE | −0.42 [−0.76, −0.08]; 0.29 | −1.39 [−1.86, −0.96]; <0.001 | −1.48 [−2.02, −1.00]; <0.001 |
| Brier | −0.06 [−0.12, −0.01]; 1.00 | −0.17 [−0.36, −0.04]; 0.004 | −0.17 [−0.38, −0.02]; 0.38 |
| NLL | −0.15 [−0.30, −0.03]; 0.11 | −0.68 [−1.55, −0.15]; <0.001 | −0.67 [−1.53, −0.14]; <0.001 |

On the held-out centre, StaRFM_bal_ and StaRFM_band_ lowered boundary ECE from 7.71% to 6.57% and 6.24%, NLL from 1.03 × 10^−2^ to 0.59 and 0.58 × 10^−2^, and the Brier score by 0.13 and 0.18 × 10^−3^ (all adjusted *p* < 0.001). Pooled ECE fell from 0.119% to 0.098% and 0.094%. These relative reductions (about 15% to 19% for boundary ECE and 43% for NLL) are of similar or larger size on the ID test set (16% to 17% and 52% to 53%). Segmentation accuracy was not harmed. On the held-out centre, StaRFM_bal_ improved DSC by 1.8 points (95% CI 0.9 to 2.6, adjusted *p* = 0.001) and StaRFM_band_ by 1.1 points (0.3 to 1.8, *p* = 0.010). HD95 differences were not significant after adjustment. On the smaller ID test set, DSC differences had wide intervals that include zero.

#### Interpretation

Five observations help to read these results.

- *Averaging matters more than the penalty itself*. Voxel-wise CMP lowered DSC at both weights tried: by 6.9 points within StaRFM at *λ*_2_ = 0.1, and by 2.1 points on its own at *λ*_2_ = 0.001, where StaRFM as a whole showed no measurable effect. The lesion-volume check is consistent with a background-dominated penalty. On the held-out centre, CMP alone shrank predicted lesions (median predicted-to-reference volume ratio 0.88 against 0.98 for the baseline; difference in log ratio −0.18, 95% CI −0.23 to −0.13).

StaRFM_bal_ left lesion volume unchanged (median ratio 0.97; difference 0.01, CI −0.02 to 0.04). Balancing the two classes, or restricting the penalty to the boundary band, turned a harmful term into one that improves every calibration metric.

- *The calibration gains come from CMP, not FIP*. FIP alone did not significantly improve any metric and slightly increased NLL on the held-out centre (+0.17× 10^−2^, CI 0.08 to 0.29). CMP_bal_ alone achieved most of the calibration gain of StaRFM_bal_ (Table 6). In this setting the contribution of FIP is not established.
- *The bounds hold but are far from tight*. All 1,968 patient-runs (eight methods, three seeds, 82 test patients) satisfied 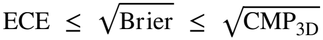. For StaRFM_bal_, mean ECE was 0.0010, against 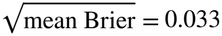 and 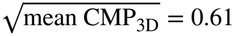. Pooled voxel ECE is small because easy background voxels dominate it, so the bounds explain the direction of the effect, not its size. Boundary ECE is the more informative quantity here. Figure 3 shows why. In the intermediate bins (mean confidence 0.55 to 0.86), every model is clearly overconfident, with accuracies between 0.47 and 0.66. Pooled ECE can nonetheless be about 0.1% only because these bins hold very few voxels: for the baseline, whose gap in these bins is at least 0.05, they can contain at most about 2% of the voxels, and nearly all the rest fall in the top bin, where predictions are almost always correct. The band variant lies closest to the diagonal in every intermediate bin on both splits, and the balanced variant in most of them, but neither removes the gap.
- *Scope*. The selected *λ*_2_ lies at the lower edge of the grid for all three forms, so smaller weights were not explored. Centre 2 was easier than the centre-1 ID test set for every method (baseline DSC 73.0% versus 59.6%), so the two splits should not be read as a measure of performance loss under shift. With only two centres, the confidence intervals reflect patient-level variability only.
- *Cost*. In a controlled profiling pass (one update on each of 20 training volumes, the first two excluded as warm-up, same GPU), StaRFM_bal_ took 12.9% longer per update than the baseline (40.2 versus 35.6 ms), and peak memory was unchanged (1.28 GB for both). Over full training, the mean epoch time rose by 9.5% (6.0 versus 5.5 s).

**Figure 3:**
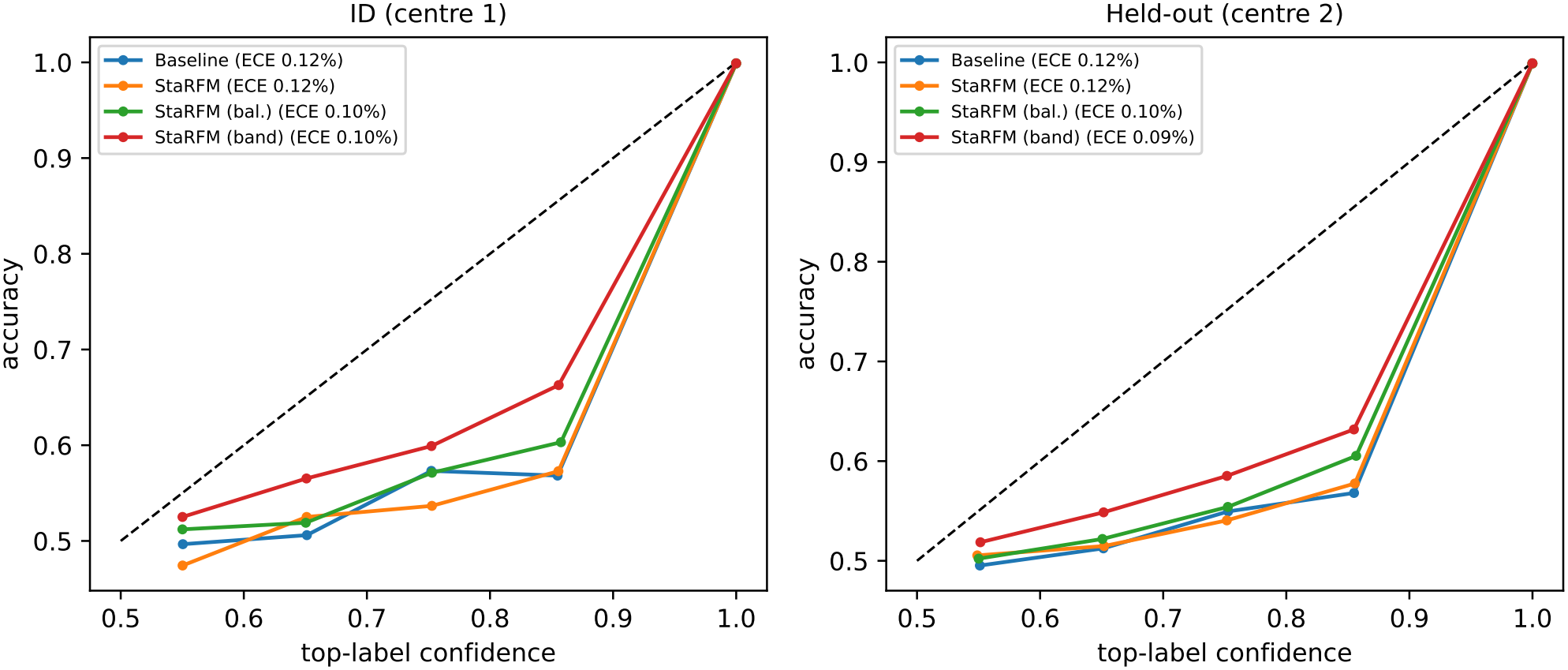
Reliability diagrams of voxel predictions in the ISLES 2022 study, for the baseline and the three StaRFM variants. Voxels are pooled over all test patients and seeds of a split (up to 200,000 sampled per patient and seed; ten equal-width bins); the legend gives the pooled ECE. Only bins that contain voxels are drawn. The dashed line marks perfect calibration. The plotted curves do not show bin occupancy; the small pooled ECE values must be interpreted alongside boundary ECE because background voxels dominate the pooled metric.

**Figure 4:**
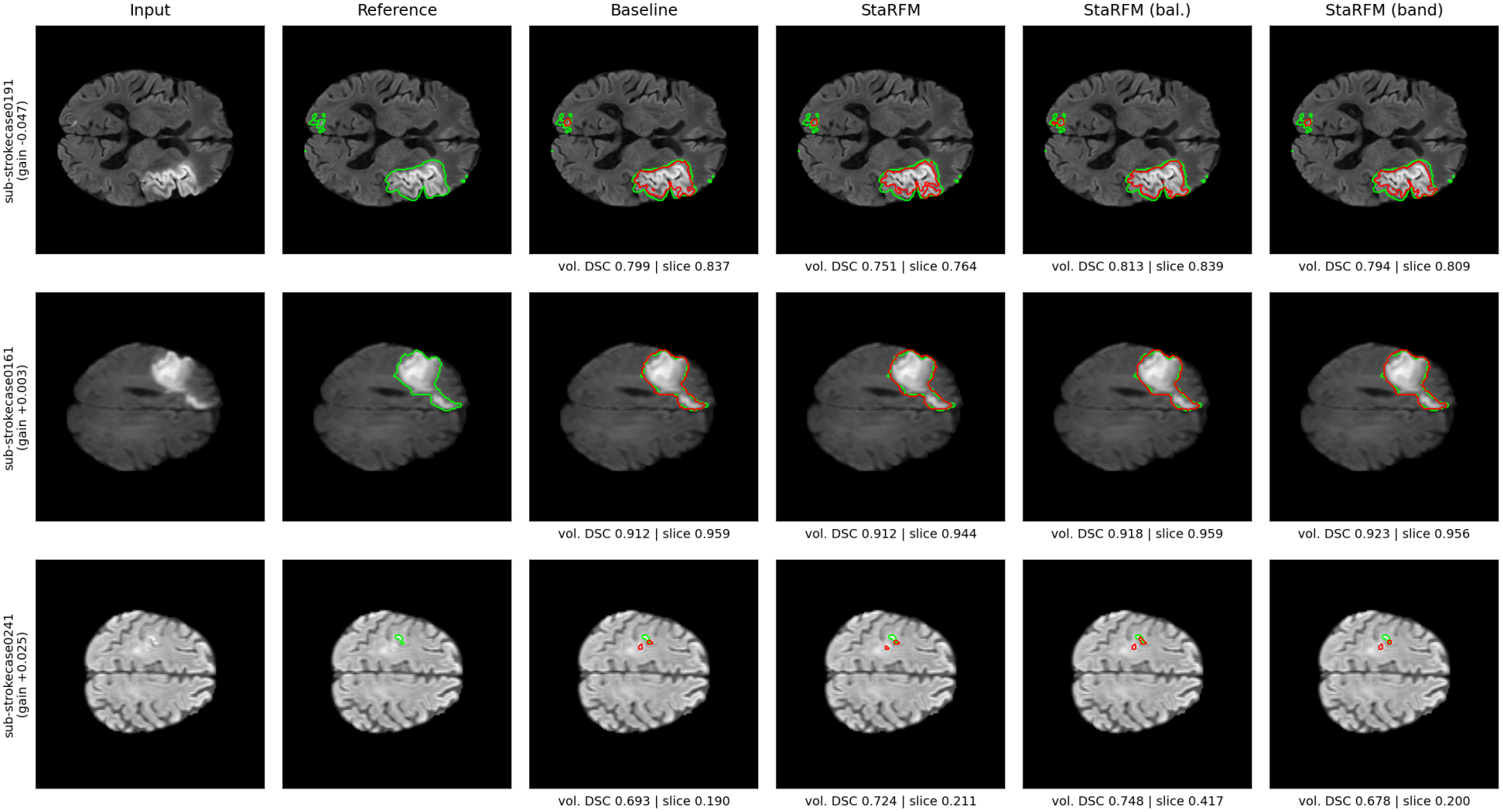
Held-out centre-2 cases from the ISLES 2022 study. Rows are the patients at the 10th, 50th and 90th percentiles of the per-patient DSC gain of voxel-wise StaRFM over the baseline (gains −0.047, +0.003 and +0.025, shown in the row labels). Each panel shows the DWI slice through the lesion centroid along the axis with the fewest voxels, with the reference (green) and the seed-mean probability map thresholded at 0.5 (red). Labels give the volumetric and slice DSC of the displayed map. The cases were chosen by a fixed rule from the test set and are illustrative only; they are not an independent validation.

### 5.5. Visual analysis

Figure 5 shows representative MRI and CT slices from the SAM experiments. SAM-FT produces spurious components near lesion boundaries, and StaRFM yields cleaner boundaries with fewer false-positive components. These images are illustrative only, and the quantitative comparison is in Table 5.

**Figure 5:**
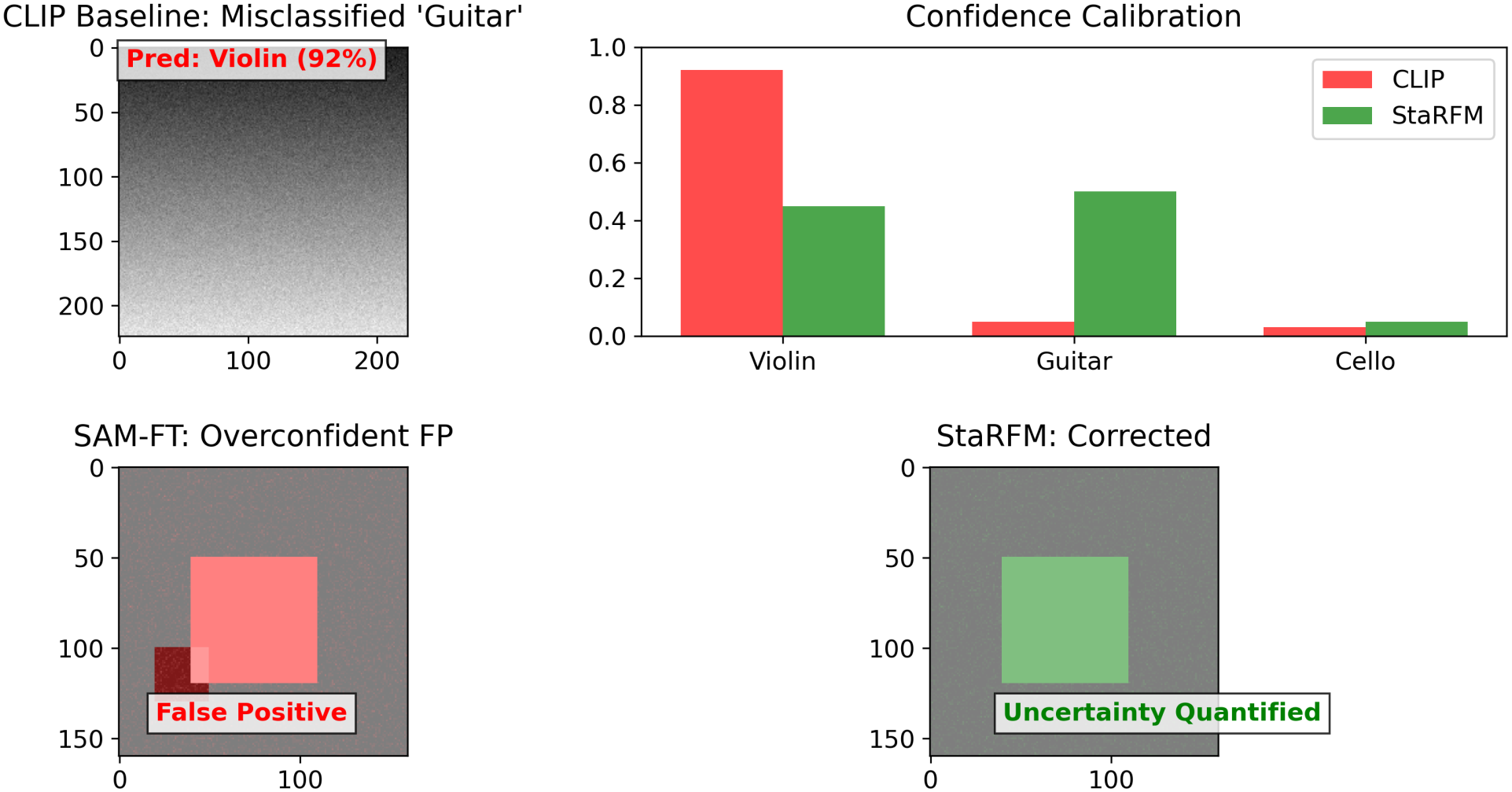
Qualitative segmentation results on representative MRI and CT slices from the SAM experiments. Columns show the input slice, the reference annotation, the SAM-FT prediction and the StaRFM prediction.

### 5.6. Ablation study

#### 5.6.1. Vision-language components

Tables 8 and 9 isolate the two penalties on the eleven recognition benchmarks. On their own, CMP raises mean accuracy from 71.3% to 72.0% and FIP to 72.5%; together they reach 73.1%. For calibration, the pattern is not what the names of the penalties suggest. CMP alone slightly *increases* mean ECE (3.17% to 3.23%), FIP alone lowers it to 3.11%, and only the combination reaches 3.03%. The two penalties are therefore complementary, and the calibration benefit comes from their interaction rather than from CMP alone. The combined model is the most accurate on nine of the eleven datasets. The exceptions are StanfordCars and EuroSAT, where every adapted variant is less accurate than CoOp.

**Table 8:**
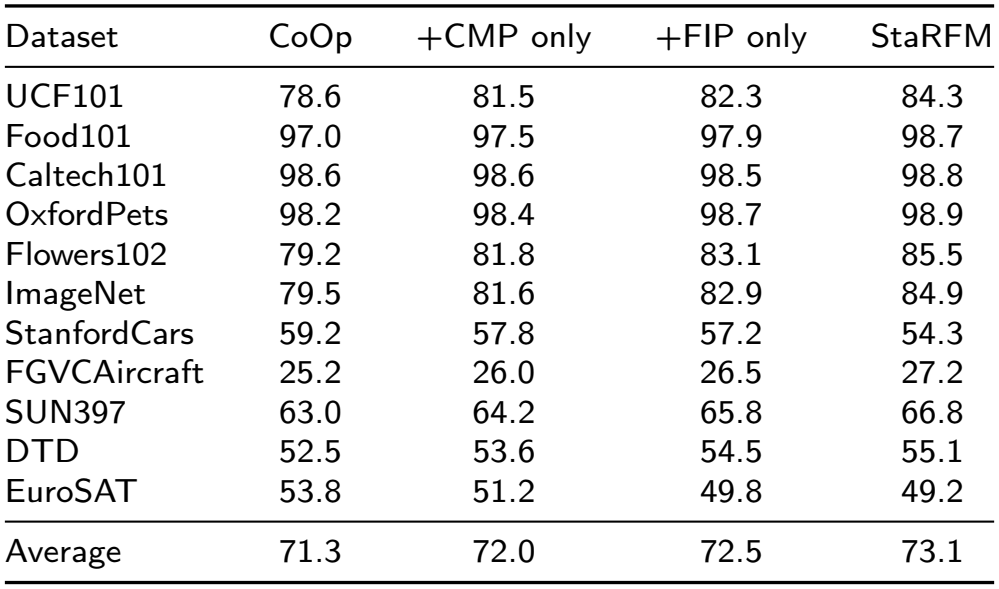
Accuracy ablation (%) on eleven vision-language benchmarks. CMP only: *λ*_1_ = 0, *λ*_2_ = 0.4; FIP only: *λ*_1_ = 0.4, *λ*_2_ = 0; StaRFM: *λ*_1_ = *λ*_2_ = 0.4. Higher is better.

| Dataset | CoOp | +CMP only | +FIP only | StaRFM |
|---|---|---|---|---|
| UCF101 | 78.6 | 81.5 | 82.3 | 84.3 |
| Food101 | 97.0 | 97.5 | 97.9 | 98.7 |
| Caltech101 | 98.6 | 98.6 | 98.5 | 98.8 |
| OxfordPets | 98.2 | 98.4 | 98.7 | 98.9 |
| Flowers102 | 79.2 | 81.8 | 83.1 | 85.5 |
| ImageNet | 79.5 | 81.6 | 82.9 | 84.9 |
| StanfordCars | 59.2 | 57.8 | 57.2 | 54.3 |
| FGVCAircraft | 25.2 | 26.0 | 26.5 | 27.2 |
| SUN397 | 63.0 | 64.2 | 65.8 | 66.8 |
| DTD | 52.5 | 53.6 | 54.5 | 55.1 |
| EuroSAT | 53.8 | 51.2 | 49.8 | 49.2 |
| Average | 71.3 | 72.0 | 72.5 | 73.1 |

**Table 9:** Calibration ablation (ECE %) on eleven vision-language bench-marks. Settings as in Table 8. Lower is better.

| Dataset | CoOp | +CMP only | +FIP only | StaRFM |
|---|---|---|---|---|
| UCF101 | 3.08 | 3.18 | 3.05 | 2.94 |
| Food101 | 3.35 | 3.42 | 3.30 | 3.02 |
| Caltech101 | 3.24 | 3.32 | 3.20 | 3.08 |
| OxfordPets | 3.06 | 3.12 | 3.00 | 2.84 |
| Flowers102 | 2.96 | 2.88 | 2.84 | 3.16 |
| ImageNet | 3.36 | 3.48 | 3.30 | 3.06 |
| StanfordCars | 3.38 | 3.50 | 3.32 | 3.16 |
| FGVCAircraft | 3.24 | 3.32 | 3.18 | 3.08 |
| SUN397 | 3.02 | 3.08 | 2.96 | 2.96 |
| DTD | 3.06 | 3.12 | 3.00 | 2.92 |
| EuroSAT | 3.08 | 3.12 | 3.02 | 3.06 |
| Average | 3.17 | 3.23 | 3.11 | 3.03 |

### 5.6.2.Medical segmentation components

Table 10 reports the component ablation on BraTS and ATLAS with the SAM backbone. Each penalty improves DSC over SAM-FT on its own, by 0.8 to 1.2 points. The combination improves it by 2.2 (BraTS) and 2.5 (ATLAS) points.

**Table 10:** Component ablation on BraTS and ATLAS (DSC %, SAM backbone). ^*^: *p* < 0.01, paired two-sided Wilcoxon signed-rank test over test patients against SAM-FT.

| Configuration | FIP | CMP | BraTS | Δ | ATLAS | Δ |
|---|---|---|---|---|---|---|
| SAM-FT |  |  | 84.0 | – | 76.8 | – |
| + FIP | ✓ |  | 85.1 | +1.1 | 78.0 | +1.2 |
| + CMP |  | ✓ | 84.9 | +0.9 | 77.6 | +0.8 |
| StaRFM | ✓ | ✓ | 86.2<sup>*</sup> | +2.2 | 79.3<sup>*</sup> | +2.5 |

#### 5.6.3. Sensitivity to the penalty weights

Tables 11 and 12 vary one weight while the other is fixed at 0.4. Mean accuracy is highest for *λ*_1_ ∈ {0.4, 0.6} (73.1% in both cases) and drops to 72.4% at *λ*_1_ = 0.8. Mean ECE changes by less than 0.2 points over *λ*_2_ ∈ [0, 0.8]. It is lowest at *λ*_2_ = 0.4 (3.03%) and rises to 3.19% at *λ*_2_ = 0.8. We use *λ*_1_ = *λ*_2_ = 0.4 for the vision-language experiments.

**Table 11:**
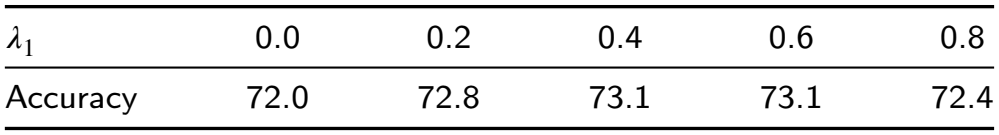
Mean accuracy (%) over the eleven benchmarks as a function of *λ*_1_, with *λ*_2_ = 0.4. Higher is better.

**Table 12:**
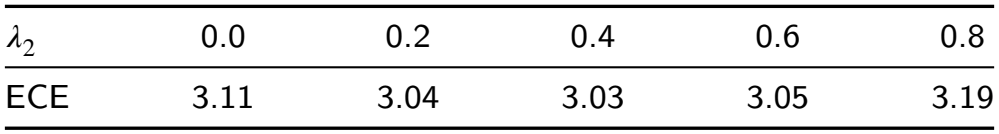
Mean ECE (%) over the eleven benchmarks as a function of *λ*_2_, with *λ*_1_ = 0.4. Lower is better.

| *λ*<sub>2</sub> | 0.0 | 0.2 | 0.4 | 0.6 | 0.8 |
|---|---|---|---|---|---|
| ECE | 3.11 | 3.04 | 3.03 | 3.05 | 3.19 |

#### 5.6.4. Computational overhead

Table 13 reports the cost of the penalties relative to CoOp. FIP requires per-sample score norms, and we compute its trace without forming the Fisher matrix. CMP operates on existing logits. Together they increase the time per epoch by 14.1% and peak memory by 8.0%.

**Table 13:** Computational overhead relative to CoOp on one NVIDIA A100 (80 GB).

| Method | Params (M) | Time/ep. (s) | Mem. (GB) | Overhead |
|---|---|---|---|---|
| CoOp | 0.03 | 142 | 11.2 | – |
| + CMP | 0.03 | 148 | 11.2 | +4.2% |
| + FIP | 0.03 | 158 | 12.1 | +11.3% |
| StaRFM | 0.03 | 162 | 12.1 | +14.1% |

### 5.7. Limitations

- **The form of CMP matters**. In the ISLES study the voxel-wise penalty, averaged over a volume dominated by background, lowered DSC at both weights tried. Its only calibration gain, a lower NLL at *λ*_2_ = 0.1, came with that loss of accuracy. Calibration gains appeared only with the class-balanced or boundary-restricted forms, which were introduced after the first round. They should be confirmed on further datasets and backbones, including the SAM experiments, which used the voxel-wise form.
- **Calibration is not guaranteed by the bounds**. Proposition 3.2 and Corollary 3.1.1 bound ECE from above, but the bounds are loose for voxel predictions. Practitioners should measure calibration on held-out sites, including near lesion boundaries. Post-hoc recalibration on a small labelled set from the deployment site is a natural complement that we did not evaluate.
- **The shift term is outside the model’s control**. The factor *D*_2_ in Proposition 3.1 depends only on the data. It is infinite when the target contains appearances that the source never produces, in which case the bound gives no guarantee.
- **Calibration metrics for the SAM experiments**. For the five-dataset SAM benchmark we report DSC only. Voxel ECE, Brier score and HD95 for these models would make the comparison complete.
- **Scope of the controlled study**. The ISLES study used a small U-Net, 30 training epochs, three seeds and two acquisition centres. The selected weights lay at the lower edge of the grid. FIP did not contribute measurably in this setting. The boundary band is defined in voxels rather than millimetres, so its physical width varies with voxel spacing.
- **Labels and hyperparameters**. Both penalties need labelled adaptation cases, and their weights may need re-tuning for datasets whose shift differs from those studied here.
- **Label shift**. The analysis assumes covariate shift. Changes in lesion prevalence or size distribution between sites are not modelled.

## 6. Conclusion

StaRFM adds a Fisher information penalty and a confidence misalignment penalty to the objectives used to adapt CLIP and SAM, and it computes both for volumetric data. Making the penalties explicit shows what they optimise. At a sigmoid head the empirical Fisher trace is a feature-weighted squared error, and voxel-wise CMP is an exponential loss on the logit margin. Making the theory precise shows what can be guaranteed. Under covariate shift, target risk is bounded by the source PAC-Bayes bound inflated by a data-dependent factor that no regulariser can shrink. Voxel ECE is bounded by the square roots of the Brier score and of CMP.

Empirically, the penalties improved accuracy on vision-language and covariate-shift benchmarks and improved Dice on five MRI and CT datasets with a SAM backbone. A controlled leave-centre-out study on ISLES 2022 qualifies these results. Averaged over all voxels, CMP lowered Dice. With lesion and background balanced, or with the penalty restricted to the boundary band, it improved boundary ECE, NLL and the Brier score on a held-out centre without harming Dice. FIP added no measurable benefit in that setting. The main practical lessons are that the averaging of a voxel-wise penalty must account for class imbalance, and that calibration must be measured on held-out acquisition sites, near lesion boundaries as well as over the whole volume.

Three directions follow. The first is to evaluate the class-balanced CMP with the SAM backbone and on more datasets and acquisition centres, with smaller penalty weights than our grid allowed. The second is to establish when, if at all, FIP adds to CMP. The third is to combine train-time penalties with post-hoc recalibration on a few labelled cases from the deployment site. None of the present results establish suitability for clinical use.

## Data availability

All datasets used in this study are publicly available from their providers: the vision-language benchmarks, BraTS, ATLAS v2.0, ISLES 2022 and the Medical Segmentation Decathlon. Code is available at https://anonymous.4open.science/r/StaRFM-8F3A/. For the study in Section 5.4, the repository contains the executable notebook, the patient split, the full configuration with the selected penalty weights, the software environment, the SHA-256 manifest of the input files, the weight-selection log, per-patient and perseed metrics, and the paired statistics under both multiple-comparison families.

## Declaration of Generative AI and AI-Assisted Technologies

During the preparation of this work the authors used ChatGPT and Grammarly in order to improve the readability and language of the manuscript. After using these tools, the authors reviewed and edited the content as needed and take full responsibility for the content of the publication.

## A. Proofs

### Proof of Proposition 3.1

i. Under covariate shift the conditional label distribution is shared, so for any non-negative loss

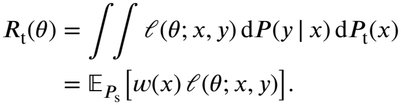 By the Cauchy-Schwarz inequality, and because 0 ≤ *l* ≤ 1 implies *l*^2^ ≤ *l*,

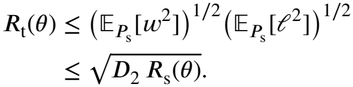 Since 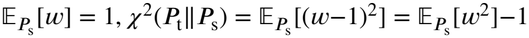, which gives *D*_2_ = 1 + *X*^2^.
ii. On the event of probability at least 1 − *δ* on which the PAC-Bayes bound holds, part (i), Jensen’s inequality for the concave square root and monotonicity of the square root give, for every *Q*,

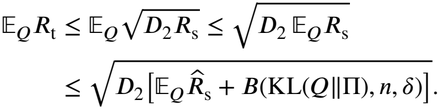 □

### Proof of Proposition 3.2

Let *q*_*v*_ = |*p*_*v*_ − *y*_*v*_ |be the probability assigned to the wrong label.

*Step 1:* (*z*_*v*_ −*c*_*v*_)^2^ = (*p*_*v*_ −*y*_*v*_)^2^ *for every voxel*. If ŷ_*v*_ = *y*_*v*_, then *z*_*v*_ = 1 and *c*_*v*_ is the probability assigned to *y*_*v*_, namely 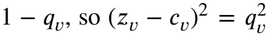. If ŷ ≠ *y*, then *z* = 0 and *c*_*v*_ is the probability assigned to the wrong label, namely *q*_*v*_, so again 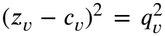. Ties 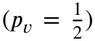 fall under one of the two cases. Hence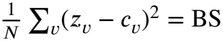.

*Step 2:* 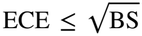. With *π*_*b*_ = |*B*_*b*_ |/*N* and |E*X*|≤ (E*X* ^2^)^½^within each bin,

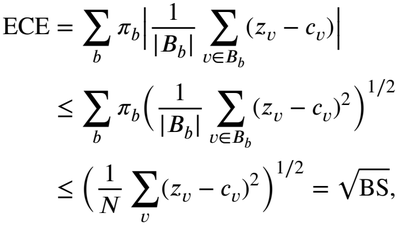

where the second inequality is Jensen’s inequality for the concave square root with weights *π*_*b*_ summing to one. Empty bins contribute nothing.

*Step 3:* BS ≤ *CMP3D*. We show 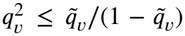. If *q* ≤ 1 − *ε*, then 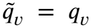 and 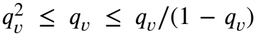. If *q* > 1 − *ε*, then 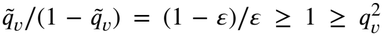 because 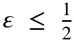. Averaging over voxels and taking square roots completes the proof. □

## Author Biographies

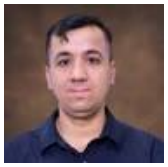

**Behraj Khan** is a Postdoctoral Researcher at St. John’s University, New York. He received his Ph.D. in Computer Science from FAST-NUCES, Karachi, Pakistan. His research interests include robust machine learning under distribution shift, Fisher information-based regularization, federated learning, and foundation model calibration. His work has appeared in venues including CVPR, ICLR, *Neurocomputing*, and *ACM Computing Surveys*.

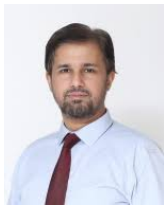

**Tahir Qasim Syed** is a Professor at the School of Computing and Computer Science, IBA Karachi, Pakistan. His research interests include machine learning, computer vision, representation learning, domain adaptation, and robust statistical learning. He has supervised doctoral and master’s research and has published extensively in international journals and conferences.

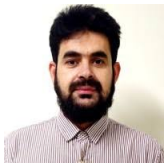

**Shabir Ahmed** is a faculty member at Gachon University, South Korea. His research interests include machine learning, data-driven modeling, intelligent systems, distribution shift, and robust learning. His research focuses on developing applied artificial intelligence methods for diverse real-world applications.

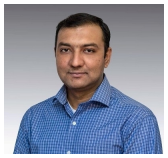

**Syed Ahmad Chan Bukhari** is a tenured Associate Professor and Director of Research at St. John’s University, New York, USA. He received his Ph.D. in Computer Science from the University of New Brunswick, Canada, and completed postdoctoral training in biomedical informatics at Yale University. His research interests include biomedical and health informatics, trustworthy artificial intelligence, machine learning, biomedical information retrieval, and semantic data integration. His research focuses on developing intelligent, data-driven methods for healthcare analytics, clinical prediction, and biomedical data interoperability.

